# Structure-based validation can drastically under-estimate error rate in proteome-wide cross-linking mass spectrometry studies

**DOI:** 10.1101/617654

**Authors:** Kumar Yugandhar, Ting-Yi Wang, Shayne D. Wierbowski, Elnur Elyar Shayhidin, Haiyuan Yu

## Abstract

Recent, rapid advances in cross-linking mass spectrometry (XL-MS) has enabled detection of novel protein-protein interactions and their structural dynamics at the proteome scale. Given the importance and scale of the novel interactions identified in these proteome-wide XL-MS studies, thorough quality assessment is critical. Almost all current XL-MS studies validate cross-links against known 3D structures of representative protein complexes. However, current structure validation approach only includes cross-links where both peptides mapped to the 3D structures. Here we provide theoretical and experimental evidence demonstrating this approach can drastically underestimate error rates for proteome-wide XL-MS datasets. Addressing current shortcomings, we propose and demonstrate a comprehensive set of four metrics, including orthogonal experimental validation to thoroughly assess quality of proteome-wide XL-MS datasets.

---

Cross-linking mass spectrometry (XL-MS) is a powerful platform capable of unveiling protein interactions and capturing their structural dynamics^1^. Although XL-MS techniques were once limited to studying individual functional complexes at a time, the development of efficient MS-cleavable chemical cross-linkers — such as disuccinimidyl sulfoxide (DSSO)^2^, disuccinimidyl dibutyric urea (DSBU)^3^ and protein interaction reporters (PIRs)^4^ — has broadened the applicability of XL-MS to proteome scale^5–8^. With the increased throughput of these techniques, the number of false positive cross-links and incorrect interactions derived from them can quickly add up with just one large-scale XL-MS experiment, if one is not careful. Therefore, thorough quality assessment methods have become critically important. Computationally defined false discovery rates (FDR) provide a quantitative metric allowing researchers to filter lists of cross-link identifications, until a theoretical quality level is achieved. In addition to FDR calculations, almost all proteome-wide XL-MS studies leverage available 3D structures of representative complexes to provide a means of validation and quality assessment^9, 10^. Here, we demonstrate fundamental flaws in this structure-based quality assessment approach that can drastically underestimate the error rates of large-scale XL-MS datasets.

The maximum theoretical distance that a given chemical cross-linker can span (e.g. 30Å for DSSO^11^) can assess agreement between existing 3D protein structures and XL-MS datasets. Particularly in small-scale studies handling intraprotein cross-links, the fraction of cross-linked residue pairs that satisfy this distance constraint may provide meaningful insights into protein flexibility or the quality of the cross-links detected. In proteome-wide XL-MS studies, researchers extend this concept and use representative, highly abundant complexes such as the ribosome and proteasome to estimate the quality of all interprotein cross-links reported. However, the majority of false positive identifications come from these interprotein crosslinks^12^. Moreover, true positive and false positive interprotein cross-links are not equally likely to successfully map onto an existing 3D structure, leading to massive underestimation of false positives (Fig. 1a). In other words, this distance-based validation effectively pre-filters the dataset, dropping most potential false positive cross-links from the evaluation entirely.

**Figure 1.**
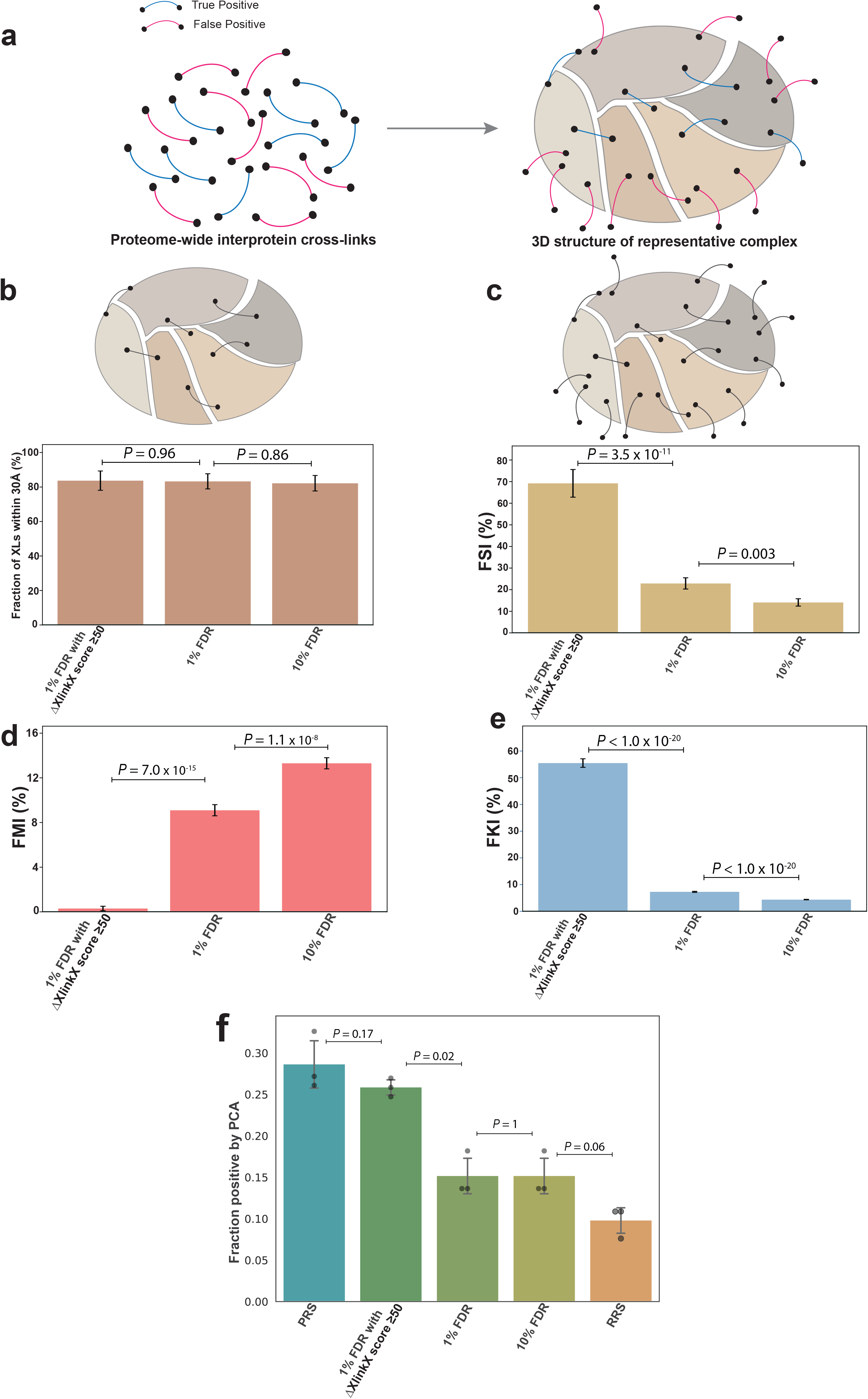
Evaluation of the conventional 3D structure-based validation approaches for proteome-wide cross-linking mass spectrometry using K562 DSSO proteome-wide XL-MS data. (a) Illustration of the limitation of current structure-mapping approach in validating the false positive XL identifications; most false positive cross-links only have one peptide mapped to the structure and are therefore ignored in the current validation approach. (b) The current structure-based validation approach is unable to distinguish datasets with drastically different quality. (c) Fraction of structure-corroborating identifications (FSI) addresses the limitation of the current approach shown in panel a. (d) Fraction of mis-identifications (FMI) differentiates the three sets in terms of their estimated percentage of mis-identifications. (e) Fraction of interprotein cross-links from known interactions (FKI) captures the underlying relative quality among the three sets. *P* values in b – e were calculated using a two-sided *Z*-test. (f) Orthogonal experimental validation of a random subset of novel interactions from the three sets using protein complementation assay (PCA). (PRS: Positive Reference Set (mean fraction positive: 0.286); RRS: Random Reference Set (mean fraction positive: 0.098); 10% FDR (mean fraction positive: 0.152); 1% FDR (mean fraction positive: 0.152); 1% FDR with ΔXlinkX score ≥50 (mean fraction positive: 0.258); The error bars represent the estimated standard error of mean; *P* values were calculated using a two-sided Welch Two Sample t-test on the log-transformed measurements; 95% confidence interval; t-statistic 4.04 for ‘10% FDR and RRS’, 7.20 for ‘1% FDR with ΔXlinkX score ≥50 and 1% FDR’, 2.13 for ‘PRS and 1% FDR with ΔXlinkX score ≥50’; 2 degrees of freedom).

Let us first assume that interprotein cross-links can be detected between any two proteins (~20,000 for human proteome-wide experiments) and that we only consider cross-links where at least one end maps to a reference protein complex structure consisting of 100 subunits. For a given cross-link, the theoretical probability that the second end also maps to this complex is ~5×10^−3^ (99/19999) and would notably be much lower for the often used ribosome (76 subunits: PDB ID 5T2C) or proteasome (34 subunits: PDB ID 5GJQ) complexes. However, these probabilities only hold for random cross-linking; cross-links derived from truly interacting proteins are much more likely to perfectly map to existing 3D structures. We expect that almost all false positive cross-links will only map one peptide to the complex structure. Current structural-mapping approaches explicitly validate cross-links where both peptides map to the same complex structure, and in doing so, they enrich for true positive cross-links and massively underestimate error rates for proteome-wide XL-MS datasets. Consequentially, these methods may erroneously annotate artifacts as novel interactions, resulting in less reliable experimental datasets for further studies.

To demonstrate our theory experimentally, we performed a proteome-wide human XL-MS experiment using the MS-cleavable cross-linker DSSO on K562 cell lysate (Methods). Next, in order to generate three sets of cross-links with drastically different qualities, we ran the XlinkX search engine (Proteome Discoverer 2.2) using three criteria of increasing stringency (10% FDR, 1% FDR and 1% FDR with ΔXlinkX score ≥50; ΔXlinkX score is a quality metric in XlinkX that allows users to filter beyond the standard FDR cutoffs). As shown in Supplementary Table 1, at 10% FDR, a set of 35,561 interprotein cross-links were identified (we intentionally chose 10% FDR to obtain a low-quality set of cross-links with many false positives), 1% FDR yielded 16,591 interprotein cross-links, whereas ‘1% FDR with ΔXlinkX score ≥50’ yielded 985 interprotein cross-links. We mapped the interprotein cross-link residue pairs from these three sets separately onto the 3D structure of the human proteasome following the conventional methodology. We then calculated the percentage of mapped residue pairs that satisfied DSSO’s theoretical constraint (≤30Å). We observed that there was no significant difference (all *P* > 0.85) among the three sets in terms of their percentage of satisfying residue pairs (Fig. 1b), even though the overall qualities of these three sets are drastically different by design. We further demonstrate this limitation of current structure-based validation by re-analyzing raw files from other publicly available datasets representing different organisms (*E. coli^6^* and Mouse11) and cellular compartments (mitochondria^11^) [Fig. 2a(i) and 2b(i)]. Our experimental results confirm that the current structure-mapping approach overestimates the fraction of true positives and thereby fails to capture the underlying error rate. Furthermore, it indicates an urgent need for additional measures to estimate the quality of proteome-wide cross-linking datasets.

**Figure 2.**
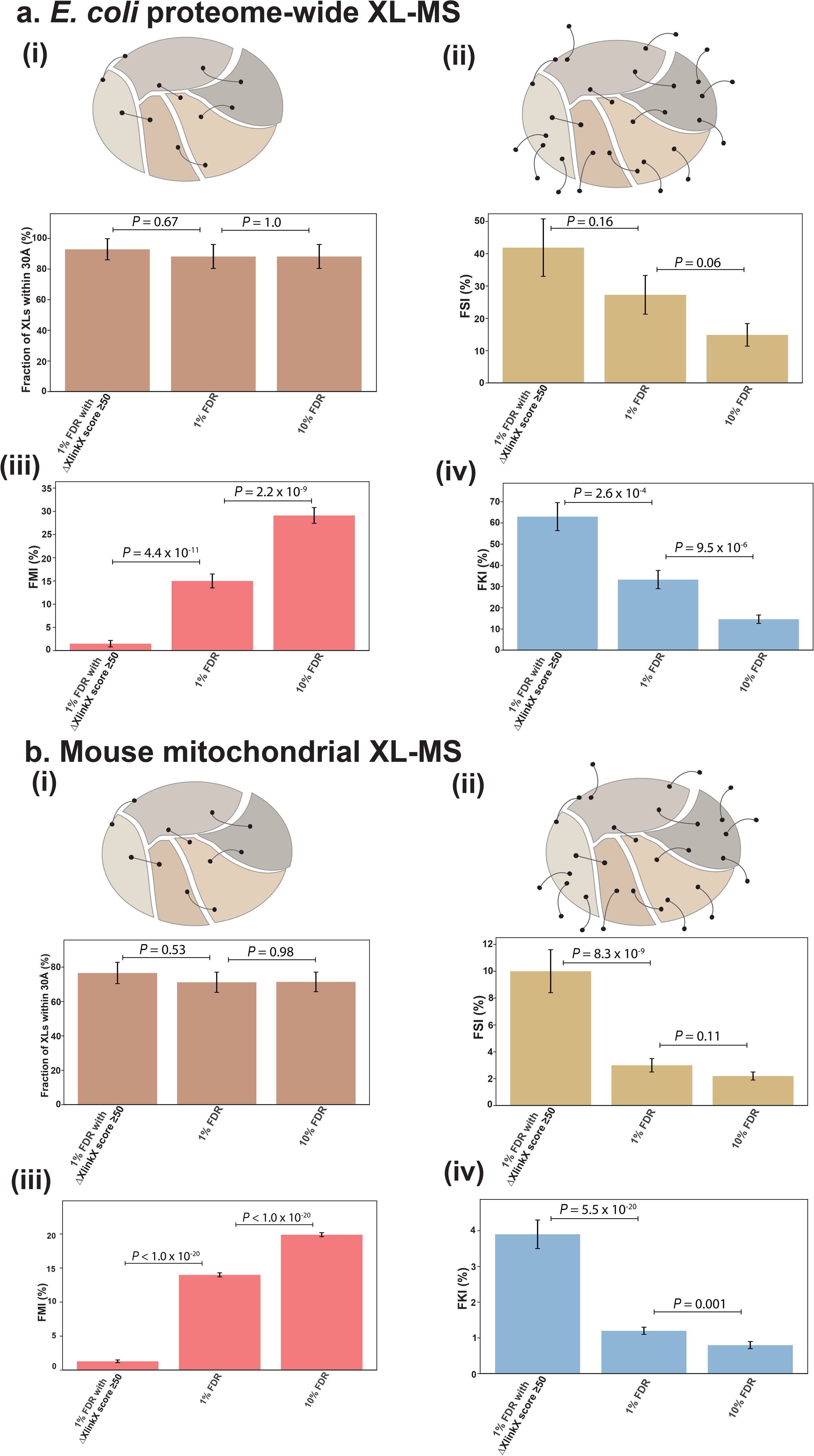
Demonstration of the utility of our comprehensive set of validation metrics on publicly available XL-MS datasets. (a) *E. coli* proteome-wide XL-MS dataset^6^: (i) Conventional structure-based validation; (ii) Fraction of structure-corroborating identifications (FSI); (iii) Fraction of mis-identifications (FMI); (iv) Fraction of interprotein cross-links from known interactions (FKI). (b) Mouse mitochondrial XL-MS dataset^11^: (i) Conventional structure-based validation; (ii) Fraction of structure-corroborating identifications (FSI); (iii) Fraction of mis-identifications (FMI); (iv) Fraction of interprotein cross-links from known interactions (FKI). *P* values were calculated using a two-sided *Z*-test; The error bars represent the estimated standard error of mean.

To address the pitfalls of current validation methods and to assess the actual quality of proteome-wide XL-MS datasets, we propose the following comprehensive set of four measurements:

i. *Fraction of structure-corroborating interprotein identifications (FSI)*: The current structure-based validation reports the fraction of cross-links supported by existing structures and consistent with the structural restrains of the linker (30Å for DSSO) relative to only those cross-links where both peptides mapped to the reference structure. As demonstrated, this removes the majority of false cross-links and reports identical fractions regardless of true underlying data quality (Fig 1a, 1b). Therefore, we propose FSI as an improved structure-based metric that uses the number of ***all*** interprotein cross-links with at least one peptide mapped to the reference structure, not just those with both peptides mapped, as the denominator (Methods).
ii. *Fraction of mis-identifications (FMI)*: Including the proteome of an unrelated organism in the search database can be an efficient way to independently assess the underlying true quality of the identified cross-links^7, 13^ (Methods). However, when choosing an unrelated organism, it is important to make sure that there is no potential experimental contamination with proteins from that organism, and homologous peptides between the two species be removed from the calculation. It should also be noted that another decoy database (reverse sequences of proteomes from both organisms) is generated for the FDR calculation in Proteome Discoverer (Methods).
iii. *Fraction of interprotein cross-links from known interactions (FKI)*: Previous studies have leveraged known protein-protein interaction networks to visualize the XL-MS datasets and infer biological insights^14, 15^. Using a metric analogous to a common machine learning term, precision^16, 17^, we leverage prior knowledge of experimentally-detected protein interactions to calculate the ‘fraction of interprotein cross-links from known interactions’ (FKI) and provide a comparative quality metric (Methods).
iv. *Fraction of validated novel interactions using orthogonal experimental assays*: Here, we propose that proteome-wide XL-MS studies need to validate a representative set of their novel interactions using an orthogonal experimental assay (e.g., Y2H, PCA), to ensure the quality and reproducibility of the dataset (Methods).

We next applied our proposed metrics on the human K562 proteome-wide XL-MS results, and demonstrated how each of them efficiently captures the differences in data quality among the three filtered sets, namely 10% FDR, 1% FDR and 1% FDR with ΔXlinkX Score ≥50 (Fig. 1). Fig. 1c shows that our improved structure-based metric FSI, differentiates the three sets with statistical significance, which could not be achieved by the conventional structure-based metric (Fig. 1b). Furthermore, Fig. 1d reveals that FMI is significantly lower at 1% FDR with ΔXlinkX score ≥50 compared to the other two sets (1% FDR has significantly lower FMI compared to that of 10% FDR). Additionally, FMI can also be corrected to account for differences in the sizes between the true and false search spaces^18^ (e.g. *S. cerevisiae* database is ~2 times larger than that of *E. coli*). The corrected FMI shows similar trend to that of uncorrected FMI across all three sets analyzed in the current study (Supplementary Fig. 1). Moreover, as shown in Fig. 1e, FKI exhibits great agreement with the expected data quality of different data sets (at 1% FDR with ΔXlinkX score ≥50, FKI is 55.5%; but at 10% FDR, FKI is merely 4.4%; *P* < 1×10^−20^). Additionally, we performed a thorough orthogonal experimental validation of randomly chosen novel interactions from the three sets using a protein complementation assay (PCA)^19, 20^. The fraction of PCA-positive novel interactions from ‘1% FDR with ΔXlinkX Score ≥50’ set (the expected highest quality set among the three sets) is distinctively higher when compared to the other two sets and indistinguishable from the positive reference set (PRS) (*P* = 0.17; Fig. 1f). Notably, the fraction of PCA-positive interactions for 1% FDR and 10% FDR are indistinguishable from that of the random reference set (RRS). We further confirmed the usefulness and robustness of the three computational metrics (namely FSI, FMI and FKI) on the re-analyzed *E. coli* (Fig. 2a) and Mouse (Fig. 2b) proteome-wide XL-MS datasets.

Taken together, our four metrics constitute a comprehensive framework to facilitate both relative comparison across different datasets and absolute estimation of error rates. Moreover, because these metrics stem from different principles, viewed individually, they provide complementary insights to various aspects of the data quality. The FMI serves as an absolute measure of error rate and should be employed as a secondary validation to the conventional FDR estimates. Because FSI typically leverages thoroughly studied complexes, in theory, it too should provide an absolute estimate of quality. Nonetheless, we do note that the metric may only provide relative comparison in cases where limited or incomplete 3D reference structures are available [Fig. 2a(ii) and 2b(ii); Methods]. The remaining two metrics namely FKI and ‘fraction of validated novel interactions’ specifically address the quality of detected interactions inferred from interprotein cross-links. The FKI can demonstrate that a given XL-MS experiment has captured physiologically relevant interactions supported by the literature. However, because a large fraction of true protein interactions is yet to be discovered, FKI can only provide relative estimates of quality among comparable datasets. Moreover, due to uneven coverage of the interactome, experiments directed at specific cellular compartments may expect unusually low FKI compared to that of cell lysates [Fig. 2b(iv)]. Though high FKI may provide general confidence that a dataset captures true interactions, explicit orthogonal experimental validation is necessary to extend this confidence to the detected novel interactions. Furthermore, using a Bayesian framework^21, 22^ (Supplementary Note 1) and leveraging the validation rates among a positive reference set (PRS) of well-known interactions and a negative reference set (RRS) of random pairs, we can calculate the absolute precision of the novel interactions detected in a XL-MS study (Supplementary Fig. 2). Especially since the true FDR at a protein pair level can be significantly higher than the estimated FDR at a peptide pair level^7, 18^, absolute precision will be critically important for confirming the quality of identified novel interactions in a cross-linking study.

Overall, we theoretically and experimentally illustrated the limitation of the current structure-based validation approach for evaluating proteome-wide XL-MS results. Additionally, we proposed a comprehensive set of four metrics, including an orthogonal experimental validation approach, and demonstrated their ability to distinguish datasets with varying qualities. Furthermore, we acknowledge that this drastic under-estimation of the error rate by structure-based measurements is unlikely to pose a serious issue for XL-MS studies on specific proteins and individual complexes as long as the cross-link search is performed against only proteins that are included in the experiment. Importantly, this issue is highly relevant for the increasingly popular proteome-wide XL-MS experiments^9, 10^ and cross-linking immunoprecipitation MS (xIP-MS) studies^23^. In these studies, the conventional structure-based evaluation (which is used in almost all studies) does not provide an adequate assessment of the data quality and will make even low-quality datasets seem good. Given the broad use and interest in these XL-MS datasets^24, 25^, the unexpected high number of false positives will severely hinder the development of the field and the utility of these datasets. Thus, it is of utmost importance to bring this problem to the attention of the field. Going forward, a comprehensive and accurate quality assessment framework such as the one proposed in this work needs to be adapted to aid in the rapid advancement of XL-MS technologies.

## ACKNOWLEDGEMENTS

We thank Rosa Viner for support in data processing with XlinkX workflow in Proteome Discoverer.

## AUTHOR CONTRIBUTIONS

H.Y. conceived and oversaw all aspects of the study. K.Y. performed the computational analyses with assistance from SDW. T.W. performed laboratory experiments with assistance from E.E.S. K.Y. wrote the manuscript with inputs from all the authors.

## Methods

### Cell culture and cell lysate preparation

The K562 cells (ATCC^®^ CCL-243™) obtained from American Type Culture Collection (ATCC) were cultured in the Iscove’s Modified Dulbecco’s Medium (IMDM) (ATCC) supplemented with 10% fetal bovine serum (FBS) (ATCC) and under humidified ambient atmosphere with 5% CO_2_ at 37°C. To prepare the cell lysate, cells were firstly collected, washed three times with cold PBS, and resuspended in cold HEPES buffer (50 mM HEPES, 150 mM NaCl, pH 7.5) supplemented with Protease Inhibitor Cocktail (Roche). The cells were then lysed by sonication on ice for 5 sec at 10% amplitude with 6 repeats. The lysate was obtained by collecting the supernatant after centrifuging the sample at 15,000 *g* for 10 min at 4°C. The protein concentration of the lysate was determined by using Bio-Rad Protein Assay Dye (Bio-Rad).

### Cross-linking reaction

The DSSO (Thermo Fisher Scientific) solution was freshly prepared by dissolving in anhydrous DMSO. The cross-linking reaction was carried out in 50 mM HEPES, 150 mM NaCl, pH 7.5 by mixing 1 mg/mL K562 cell lysate with 1 mM DSSO for 1 hour at room temperature. The reaction was terminated by 50 mM Tris-Cl buffer, pH 7.5.

### Cross-linked sample processing for analysis

The DSSO-cross-linked lysate was denatured in 1% sodium dodecyl sulfate (SDS) at 65°C for 15 min. The lysate was then reduced by dithiothreitol (DTT) and alkylated with iodoacetamide. The proteins were extracted and precipitated by mixing the sample with excessive cold acetone-ethanol solvent (acetone:ethanol:acetic acid=50:49.9:0.1, v/v/v). The precipitates were collected and digested by TPCK-treated trypsin (Worthington Biochemical Corporation) in 50 mM Tris-Cl, 150 mM NaCl, 2 M Urea, pH 8.0 at 37°C overnight. After trypsinization, the sample was acidified by 2% trifluoroacetic acid-formic acid (TFA-FA) solution and desalted through Sep-Pak C18 cartridge (Waters). The collected elutes were dried using the SpeedVac^TM^ Concentrator (Thermo Fisher Scientific) and stored at −80°C for further analysis.

### Fractionation of cross-linked peptides by Strong Cation Exchange (SCX)

The 1 mg of desalted trypsin-digested sample was reconstituted in 25% acetonitrile/0.1% formic acid (v/v) and passed through a Spin-X centrifuge tube filters (Corning). The filtered sample was then fractionated by HPLC using a SCX column: PolySULFOETHYL A column (5 µm, 200 Å, 2.1 × 200 mm; PolyLC) operated by Dionex UltiMate 3000 Series HPLC (Thermo Fisher Scientific). Two following buffers were used: 10 mM potassium phosphate monobasic in 25% acetonitrile, pH 3.0 (Buffer A) and 10 mM potassium phosphate monobasic plus 500 mM potassium chloride in 25% acetonitrile, pH 3.0 (Buffer B). The fractionation was performed using a linear gradient of 5-60% of Buffer B in 40 min with an additional 60-100% of Buffer B in 10 min at a flow rate of 200 µL/min. A total of 60 fractions were collected at 1-min intervals. The fractions obtained from 23 to 60 min were pooled into 25 fractions and desalted individually using SOLA HRP SPE cartridges (Thermo Fisher Scientific). The eluted peptides were dried by SpeedVac^TM^ Concentrator (Thermo Fisher Scientific) and stored at −20°C for LC-MSn analysis.

### Fractionation of cross-linked peptides by hydrophilic interaction liquid chromatography (HILIC)

The desalted and dried sample was dissolved in 70% acetonitrile and 1% formic acid. The cross-linked peptides were then enriched by HILIC on the Dionex UltiMate 3000 Series HPLC instrument (Thermo Fisher Scientific) equipped with a TSKgel Amide-80 column (3 µm, 4.6 mm × 15 cm; Tosoh). Solvent A containing 90% acetonitrile, solvent B composed of 80% acetonitrile and 0.005% trifluoroacetic acid, and solvent C comprising 0.025% trifluoroacetic acid were used. The fractionations were performed using the following gradients: 0-5 min (0-98% B and 0-2% C); 5-55 min (98-75% B and 2-25% C); and 55-60 min (75-5% B and 25-95% C) at a flow rate of 600 600 µL/min. The fractions were collected from 5-55 min with 30-sec intervals. Each fractions were dried individually by SpeedVac.

### LC-MS^n^ analysis

The SCX fractionated samples were analyzed on an Acclaim PepMap C18 nano column (3 µm, 75 µm × 25 cm, Thermo Fisher Scientific) using UltiMate3000 RSLCnano (Dionex) coupled to an Orbitrap Fusion (Thermo Fisher Scientific) mass spectrometer. The chromatographic separation was achieved by 5-40% B at 300 nL/min in 120 min. The CID-MS2-MS3 workflow was used for MS^n^ data acquisition. The MS1 precursors were detected in Orbitrap mass analyzer at m/z = 375-1575 and resolution = 60,000. The precursor ions with the charge of 4+ to 10+ were selected for CID-MS2 acquisition in Orbitrap mass analyzer at resolution = 30,000, AGC target = 5 × 10^4^, isolation width = 1.6 m/z, maximum injection time = 100 ms, and collision energy = 25% of CID. The peaks captured in the CID-MS2 spectrum with a mass difference of 31.9721 Da triggered the following acquisition of CID-MS3 spectra in Ion Trap with the collision energy of CID at 35% and AGC target at 2 ×10^4^. All spectra were recorded and analyzed using Xcalibur 3.0 software and Orbitrap Fusion Tune Application v. 2.1 (Thermo Fisher Scientific).

The HILIC fractions were analyzed using an EASY-nLC 1200 system (Thermo Fisher Scientific) equipped with an in-house packed capillary column (125-µm × 25-cm) and coupled online to an Orbitrap Fusion Lumos Tribrid mass spectrometer (Thermo Fisher Scientific). The column was packed with 3-µm C18 resin (Michrom BioResources). The LC analysis was performed using 10−40% B for 180 min at 300 nl/min with solvent A composed of 0.1% formic acid (FA) and solvent B composed of 80% acetonitrile (ACN) and 0.1% FA. For MS^n^ data acquisition, the CID-MS2-HCD-MS3 method was used. The MS1 precursors were detected in Orbitrap mass analyzer at m/z = 375-1500 and resolution = 60,000. The precursor ions with the charge of 4+ to 8+ were selected for MS2 analysis in Orbitrap mass analyzer at resolution = 30,000, AGC target = 1 × 105, isolation width = 1.6 m/z, maximum injection time = 10^5^ ms, and collision energy of CID at 25%. The peaks with a mass difference of 31.9721 Da in MS2 spectra were selected for further MS3 analysis. The selected ions were fragmented in Ion Trap using HCD with the collision energy at 35% and AGC target of 2 × 10^4^. All spectra were recorded by Xcalibur 4.1 software and Orbitrap Fusion Lumos Tune Application v. 3.0 (Thermo Fisher Scientific).

### Data processing

Cross-links were identified using XlinkX software (Proteome discoverer 2.2). PD templates for different XlinkX search methodologies were obtained from Rosa Viner (Thermo fisher Scientific). Target sequences were downloaded from Uniprot database^26^ (with filter ‘reviewed’): ((i) *Escherichia coli*: 5268 sequences; downloaded on 28^th^ October 2017, (ii) *Saccharomyces cerevisiae:* 7904 sequences; downloaded on 28^th^ September 2017, (iii) Human (*Homo sapiens)*: 42202 sequences; downloaded on 23^rd^ June 2017), and (iii) Mouse (*Mus musculus)*: 17019 sequences; downloaded on 8^th^ July 2019). When performing searches for Fig. 1d, XlinkX crashed multiple times given the huge number of raw files (122 files) and the enormous search space (*H. sapiens* + *S. cerevisiae*). Hence, we ran the searches on a smaller set of raw files (25 files) to generate Fig. 1d.

### Mapping of cross-links to existing PDB structures

Cross-links from our K562 proteome-wide XL-MS dataset were mapped to the 3D structure of human 26S proteasome structure (PDB id: 5GJQ) utilizing residue level mappings between Uniport and PDB entries obtained from SIFTS^27^ database. For the re-analyzed mouse mitochondrial dataset^11^, existing representative complexes for the host species were not available. Therefore, the cross-links were instead mapped to homologous complexes (PDB IDs 1EUC, 1T9G, 5LNK, 1ZOY, 1NTM, 1V54) as shown previously^11^. In brief the protein sequences for all proteins involved in detected cross-links were aligned against a reference database containing PDB sequences of interest using BLAST^28^. All BLAST matches with significant E-value and percent identity greater than 70% were retained. Exact positions for each cross-link were mapped against homologous PDB structures using a pairwise alignment, and cross-links were only considered successfully mapped if the cross-linked lysine was conserved in the structure. In cases where multiple positions within the PDB structure were valid, the mapping that produced the shortest length cross-link was prioritized. Any cross-links where the exact position of the cross-linked lysine was not structurally resolved in a homologous PDB structure were considered partially mapped. We utilized the above-mentioned homology-based approach for E. coli dataset^6^ due to the unavailability of SIFTS residue-level mapping for most of the representative structures (PDB IDs 2VRH, 1DKG, 1PCQ, 3JCD, 4PC1, and 2LRX).

### Fraction of structure-corroborating identifications (FSI)

FSI can be calculated using the following equation:

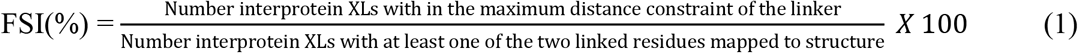

In this work, we used 30Å as the maximum distance constraint for DSSO.

### Fraction of mis-identifications (FMI)

FMI is the fraction of cross-link identifications from a false search space (from an unrelated organism) among all the identified cross-links. It can be calculated using the following equation:

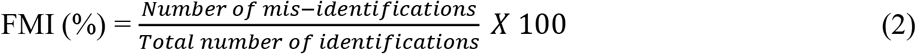

All the raw files were re-analyzed against a sequence database containing all the sequences from the target organism’s proteome and all the sequences from *S. cerevisiae* proteome. Then the fraction of cross-links with at least one of the two linked residues unambiguously mapped to proteins from *S. cerevisiae* is calculated (if any cross-link has peptide shared between homologous proteins from the target organism and *S. cerevisiae*, it was considered a true identification). It is important to note that FMI is estimated after the cross-link results are filtered at a conventional FDR threshold (1% FDR in the current study). It also noteworthy that like the conventional FDR calculations^29^, FMI calculations can also be sensitive to drastic differences in sizes of the proteome database of the unrelated organism. We utilized the following equation adapted from Fischer and Rappsilber^18^ to account for difference in database sizes.

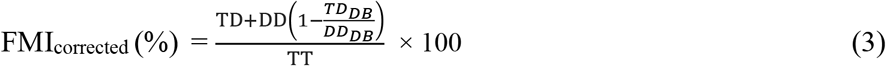

where, TT is the number of target-target matches, DD is the number of decoy-decoy matches, and TD is number of target-decoy and decoy-target matches. TD_DB_ is the number of all possible unique target-decoy and decoy-target peptide pairs and DD_DB_ is the number of all possible unique decoy-decoy peptide pairs.

### Fraction of interprotein cross-links from known interactions (FKI)

FKI for proteome-wide XL-MS studies can be defined as the fraction of the identified interprotein cross-links from previously known protein-protein interactions. It can be derived using the following equation:

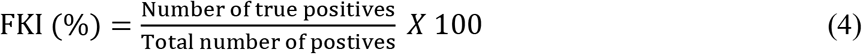

where, “positives” refer to all the identified interprotein cross-links, and “true positives” refer to interprotein cross-links from known protein-protein interactions. We compiled the known protein-protein interactions for *E. coli* (24,745), Mouse (40,527) and Human (336,033) from seven primary interaction databases. These databases include IMEx^30^ partners IntAct^31^, MINT^32^, and DIP^33^; IMEx observer BioGRID^34^; and additional sources HPRD^35^, MIPS^36^, and iRefWeb^37^. Furthermore, iRefWeb combines interaction data from CORUM^38^, BIND^39^, MPPI^36^ and OPHID^40^. We converted all gene identifiers in each database to Entrez gene IDs and then mapped to uniport IDs.

### Fraction of validated novel interactions using orthogonal experiment, namely protein complementation assay (PCA)

The ORFs of novel protein-protein interactions in pDONR223 vector was cherry-picked from hORFeome v8.1 library^41^. In each of the categories, namely 1% FDR with ΔXlinkX Score ≥50, 1% FDR, and 10% FDR, 93 protein pairs were randomly picked without any overlaps between categories. The Gateway LR reactions were performed to clone the individual bait and prey protein of each protein pair into the expression plasmids containing the complementation fragments of the fluorescent protein Venus. To perform the assay, the HEK293T cells were prepared in Dulbecco’s Modified Eagle Medium (DMEM) supplemented with 10% fetal bovine serum (FBS) (ATCC) in black 96-well flat-bottom plates (Costar) with 5% CO_2_ at 37°C. When reaching 60-70% confluency, the cells were co-transfected with both plasmids containing the Venus fragments-tagged bait and prey ORF (100 ng for each) which were pre-mixed and incubated with polyethylenimine (PEI) (Polysciences Inc.) and OptiMEM (Gibco). For positive and negative controls, the set containing previously published 92 positive reference pairs and 92 negative reference pairs were simultaneously examined^20, 42^. After 58 hours, the fluorescence intensity of the transfected cells was measured and recorded using Infinite M1000 microplate reader (Tecan) (excitation = 514 ± 5 nm / emission = 527 ± 5 nm). The PCA experiments were performed and analyzed in triplicate.

### Statistics

Statistical analyses were performed as indicated in the figure legends. Exact *P* values are provided for all compared groups.

### Data availability

Raw files from the human proteome XL-MS experiment and any additional data that supports the findings of this study are available from the corresponding author upon request.

