## Supplementary Figures for "Structure-based validation can drastically under-estimate error rate in proteome-wide cross-linking mass spectrometry studies"

**Supplementary Figure 1**. Corrected FMI for the three datasets analyzed in the study (Utilizing equation 3 in Methods section). (a) Human proteome XL-MS, (b) *E. coli* proteome XL-MS and (c) Mouse mitochondrial XL-MS.

**Supplementary Figure 2**. Estimated precision using PCA experiments for the three datasets of different quality from the human K562 proteome-wide XL-MS experiment. See Supplementary Note 1 for a detailed description of the methodology.

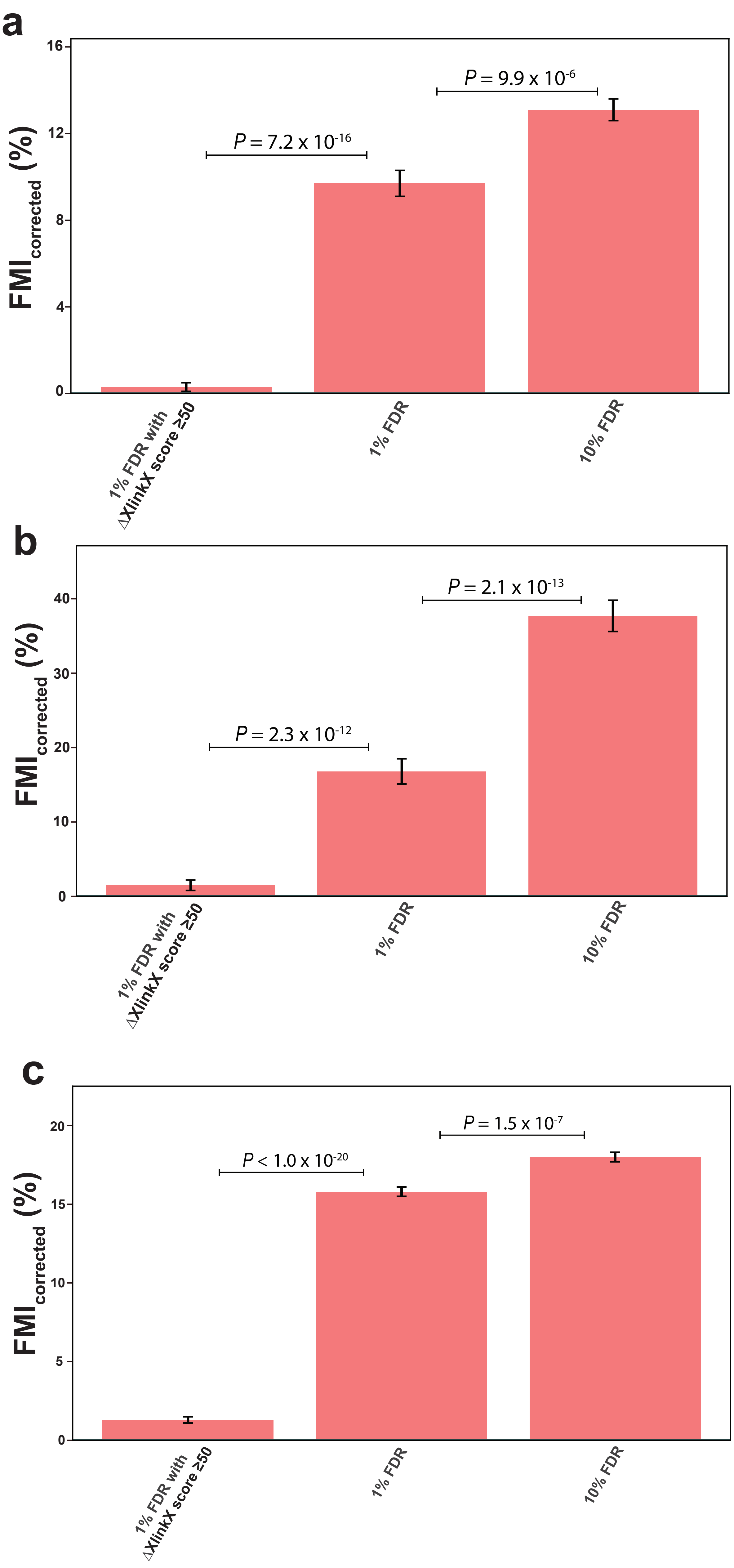


**Supplementary Figure 1**

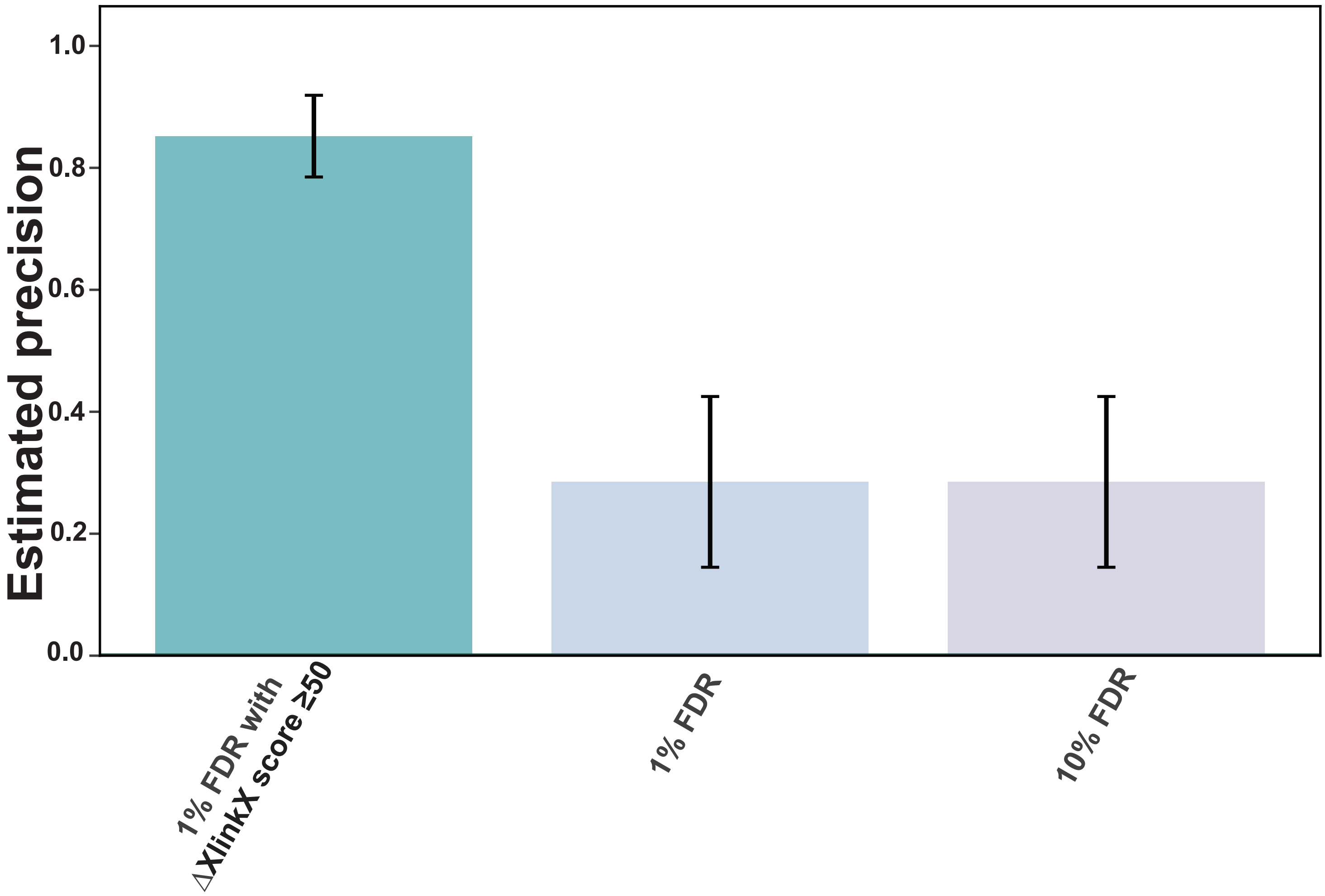


Supplementary Figure 2
