## Supplementary Note for "Structure-based validation can drastically under-estimate error rate in proteome-wide cross-linking mass spectrometry studies"

**Supplementary Note 1:** Experimental estimation of precision and its standard error for different XL-MS sets using PCA assay

As described before[1, 2], we can estimate the underlying absolute precision of a given cross-linking mass spectrometry (XL-MS) dataset using results from our PCA experiments. Briefly, using Bayes' rule, we can build relationships between true and false positive rates from different XL-MS sets and observed positive interactions by PCA as follows:

$$\Pr(A+|Y+) = \Pr(A+|Y+,T+) \times \Pr(T+|Y+) + \Pr(A+|Y+,T-) \times \Pr(T-|Y+)$$

where A+ correspond to observing a positive interaction in PCA, Y+ corresponds to observing a positive interaction in a given XL-MS dataset, and T+(T-) corresponds to an interaction being a real positive (negative) interaction. The precision of XL-MS set is the term Pr(T + | Y +) [which is also equal to 1-Pr(T - | Y +)].

Assuming conditional independence between XL-MS and PCA, we can write:

$$\Pr(A+|Y+) = \Pr(A+|T+) \times \Pr(T+|Y+) + \Pr(A+|T-) \times \Pr(T-|Y+)$$

Solving for the precision of the XL-MS sets yield:

$$\Pr(T+|Y+) = \frac{\Pr(A+|Y+) - \Pr(A+|T-)}{\Pr(A+|T+) - \Pr(A+|T-)}$$

Pr(A + | T +) and Pr(A + | T- ) were measured in the positive reference set (PRS) and random reference set (RRS) experiments. So, for the different XL-MS sets, we can calculate precision as:

$$Precision = \frac{F_{XL-MS} - F_{RRS}}{F_{PRS} - F_{RRS}}$$

Where, $F_{XL\text{-}MS}$ is the fraction positive by PCA for a random subset of novel interactions from a given XL-MS dataset, which is the best estimator for Pr(A + | Y +); $F_{PRS}$ is the fraction positive by PCA for PRS, the estimator for Pr(A + | T +); and $F_{RRS}$ is the fraction positive by PCA for RRS, the estimator for Pr(A + | T -).

To estimate the standard error of the precision, we used the standard delta method:

$$\sigma_X^2 = (\frac{\partial f}{\partial A}\sigma_A)^2 + (\frac{\partial f}{\partial B}\sigma_B)^2 + (\frac{\partial f}{\partial C}\sigma_C)^2$$

where, *X* = f(*A*, *B*, *C*, …). *A*, *B*, *C*, … are independent random variables.

Here, the standard error of the precision is calculated as:

$$\sigma_{precision} = \sqrt{(\frac{1}{F_{PRS} - F_{RRS}})^2 \times \sigma_{XL-MS}^2 + (\frac{F_{XL-MS} - F_{RRS}}{F_{PRS} - F_{RRS}})^2 \times \sigma_{PRS}^2 + (\frac{F_{XL-MS} - F_{PRS}}{F_{PRS} - F_{RRS}})^2 \times \sigma_{RRS}^2}$$
